## Supplemental Information for "A method for extracting effective interactions from Hi-C data with applications to interphase chromosomes and inverted nuclei"

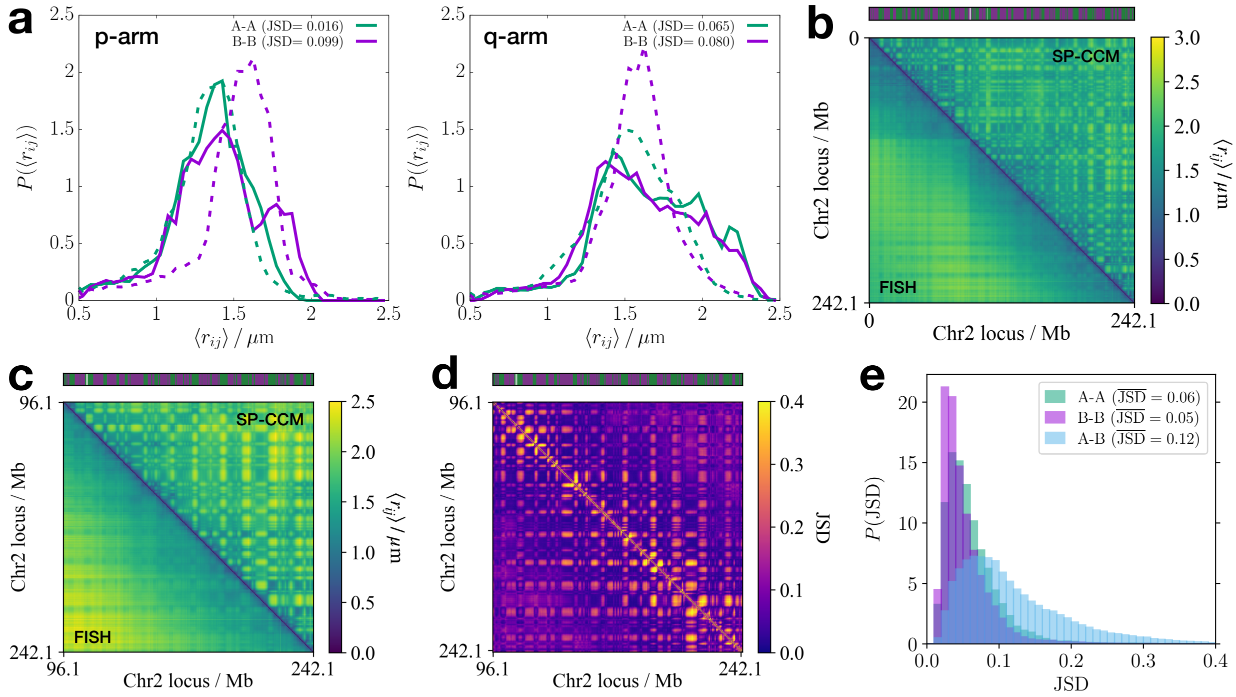


FIG. S1. Comparison between Chr2 structures from FISH experiments [43] and SP-CCM polymer simulations. (a) Plots of the probability distributions of the mean pair distance, $\langle r_{ij} \rangle$, computed separately for the p- (left) and q-arms (right) of IMR90 Chr2. Results from experiments [43] and simulations are shown in solid and dashed lines, respectively. The simulation length unit was converted to the real scale using $\sigma = 0.2\,\mu$m, and the distributions for A-A and B-B pairs are shown in green and purple colors, respectively. The small JSD values between experiments and simulations are small, which shows excellent agreement. (b) Comparison between the mean pairwise distance matrices for the whole Chr2, obtained from imaging experiments (lower triangle) and SP-CCM simulations (upper triangle). (c) Same as panel b except that the data are shown for the q-arm only. (d) Heatmap of the JSD matrix, where each element indicates the value of JSD between the probability distributions, $P(r_{ij})$, for a given locus pair, which are computed from the imaging experiments and the simulations. (e) Histograms of JSD shown in panel d for a given type of locus pair. The JSD values are small, which implies excellent agreement between simulations and experiments.

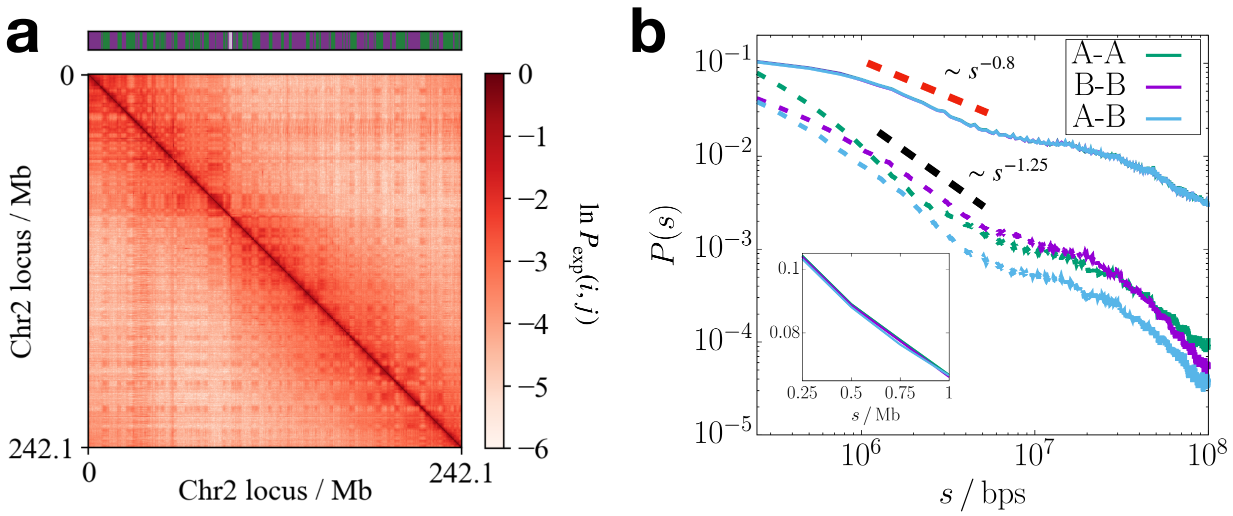


FIG. S2. Difference in the structure ensembles between the FISH and the Hi-C data. (a) CM for the IMR90 Chr2 calculated using the 3-D-structures obtained from the imaging FISH experiment. (b) Contact probability at a given genomic distance, $P(s)$, computed for different pair types. The solid and dashed lines show the results for the FISH and the Hi-C experiments, respectively. The inset shows the magnified view of $P(s)$ for the FISH data. Note that there is only negligible difference in $P(s)$ between A-A (or B-B) and A-B pairs for the FISH data in contrast with $P(s)$ for the Hi-C data. In addition, the power exponent in $s$ is different between the FISH and the Hi-C data, as shown by the red and black dashed lines.

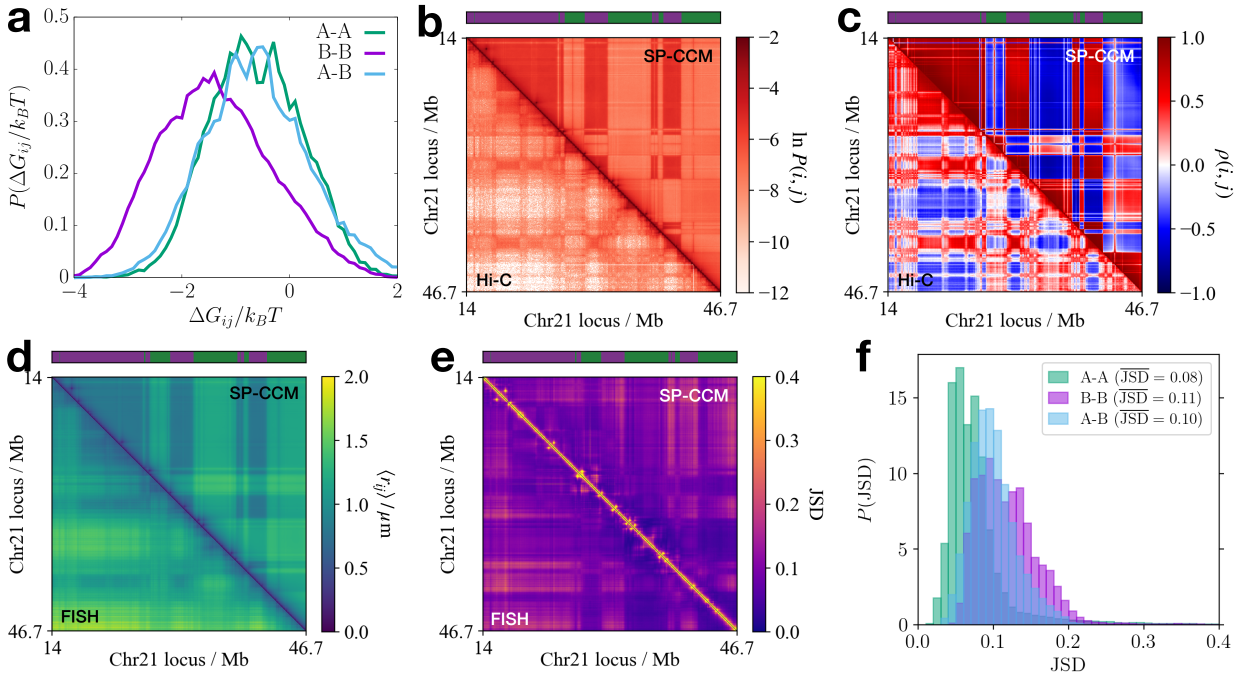


FIG. S3. Comparison between the results for Chr21 from experiments and simulations. (a) Distributions of the SP interaction energies for A-A, B-B, and A-B locus pairs, shown in green, purple, and sky-blue colors, respectively. (b) Comparison between the contact matrices obtained from the Hi-C experiment (lower triangle) and the SP-CCM simulation (upper triangle). The bar above the map shows the A/B compartment type (green/purple) of the individual loci in Chr21. (c) Pearson correlation matrices computed from the contact matrices shown in panel b. (d) Comparison between the mean pairwise distance matrices, $\langle r_{ij} \rangle$, obtained from the imaging experiments and the simulations. (e) JSD matrices, where each element indicates the value of the JSD between the probability distributions, $P(r_{ij})$, for a given locus pair computed from the imaging experiments and the simulations. (f) Histograms of the JSDs shown in panel e for a given type of loci pair.

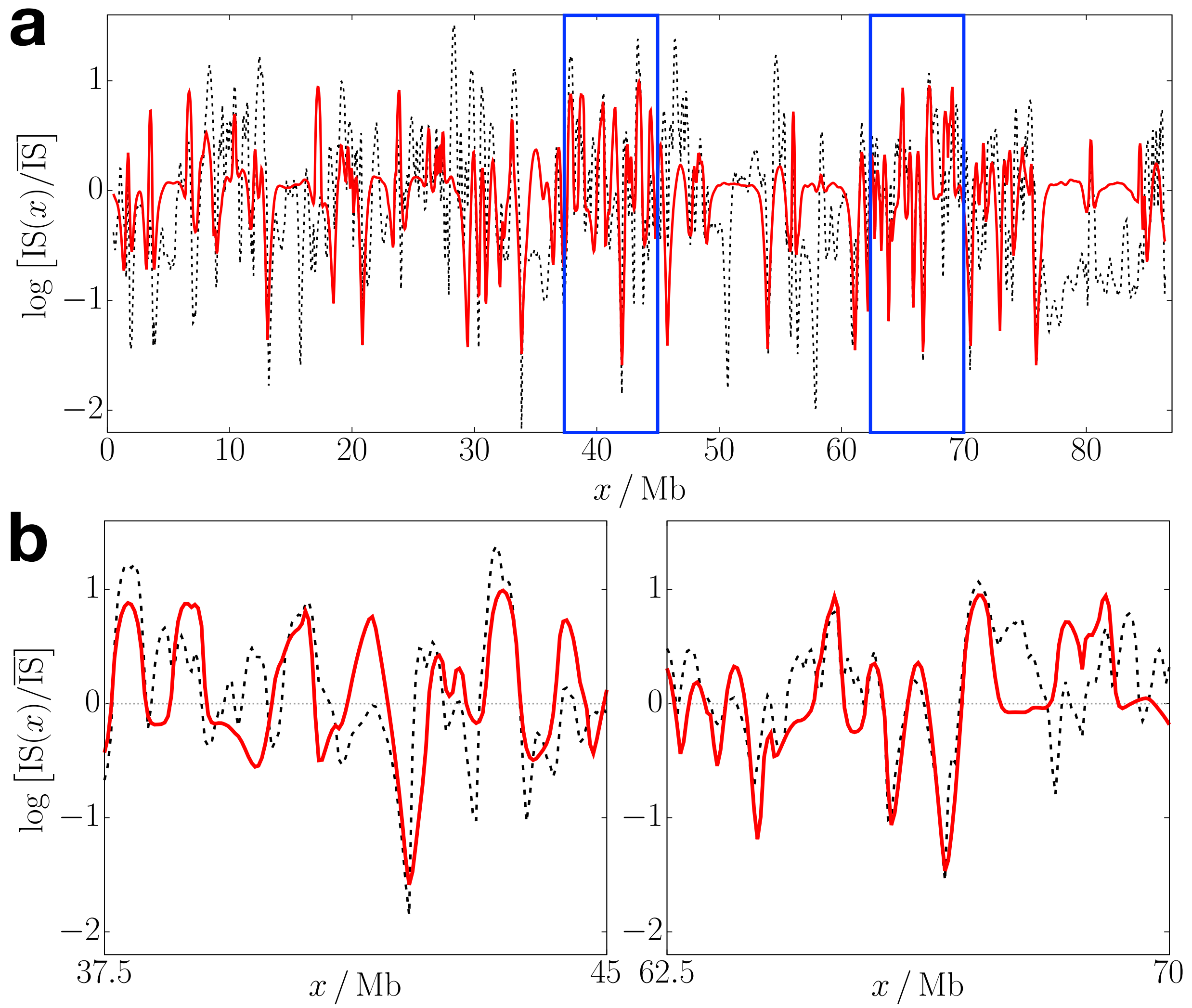


FIG. S4. Comparison between the insulation profiles, $\log\left[\mathrm{IS}(x)/\overline{\mathrm{IS}}\,\right]$, calculated using the SP-CCM simulated CM (solid line) and the Hi-C inferred CM (dashed line); $x$ denotes the position of the locus of Chr2 and $\overline{\mathrm{IS}}$ is the mean value of $\mathrm{IS}(x)$ over the p-arm. Panel a shows the plots over the whole region of the simulated p-arm. Panel b shows the magnified views of the blue-boxed regions in panel a. The corresponding contact maps are shown in Fig. 2A of the main text.

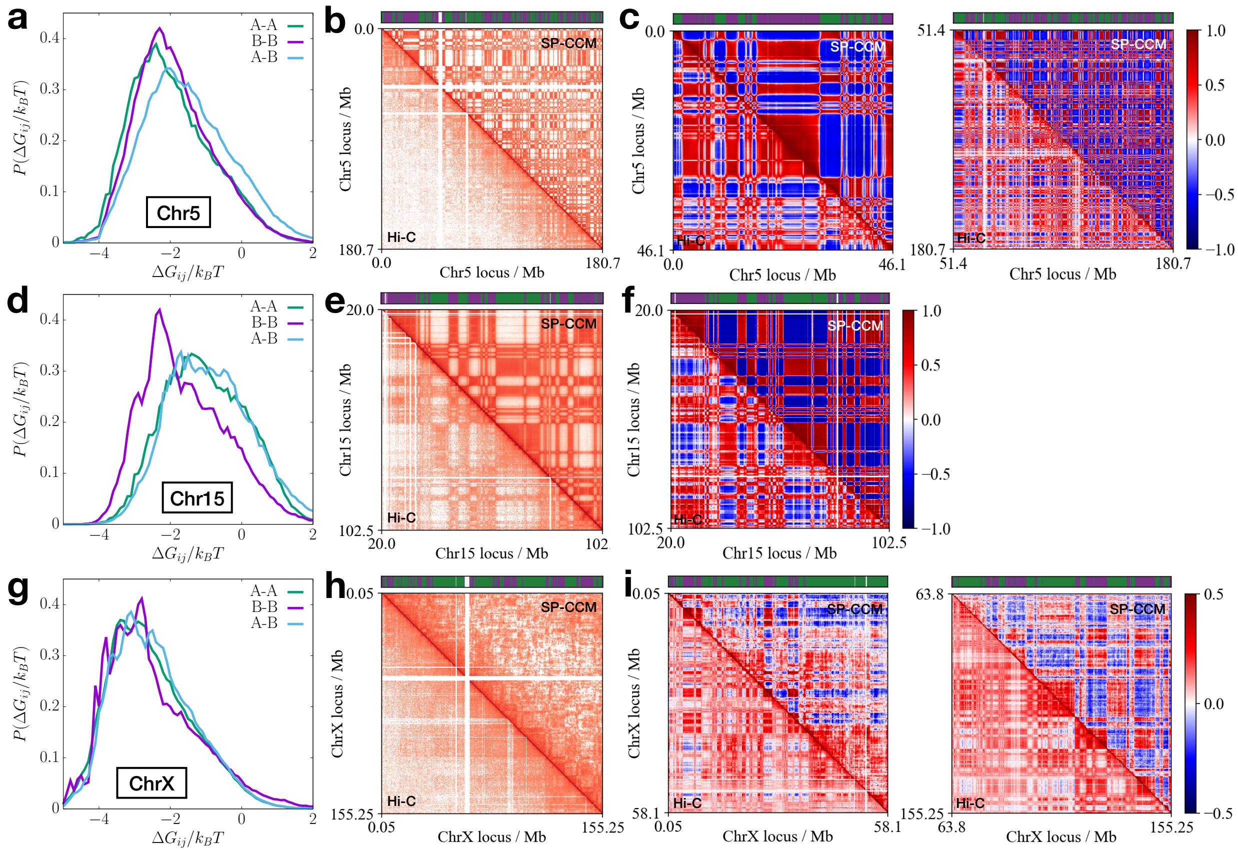


FIG. S5. SP-CCM simulation results for Chr5, CHr15, and ChrX from the IMR90 cell line. (a,d,g) Distributions of the interaction energies for A-A, B-B, and A-B loci pairs, shown in green, purple, and sky-blue colors, respectively. (b,e,h) Comparison between the contact matrices obtained from the Hi-C experiment (lower triangle) and the SP-CCM simulations (upper triangle). The bar above the map indicates the A/B compartment type (green/purple) of the individual loci. (c,f,i) Pearson correlation matrices computed from the contact matrices. Note that panels c and i show the results for p (left) and q (right) arms separately, whereas panel f shows the result for q-arm only (p-arm is not sequenced).

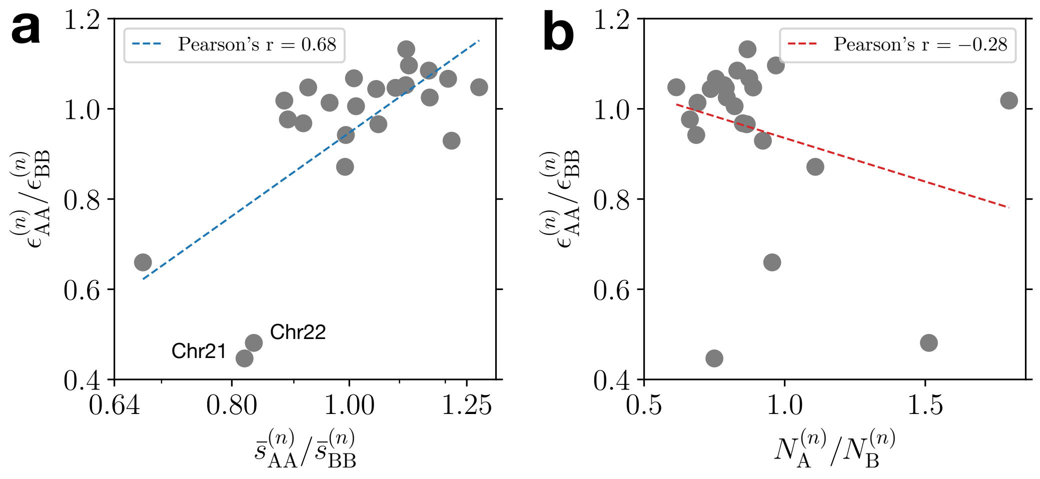


FIG. S6. Relationship between the mean SP value and chromosome sequence for IMR90. (a) Scatter plot of the ratio between $\epsilon_{\mathrm{AA}}$ and $\epsilon_{\mathrm{BB}}$ for each chromosome with respect to the ratio of average genomic distance for A-A pairs to that for B-B pairs. The $x$-axis is in the log scale. (b) Scatter plot of $\epsilon_{\mathrm{AA}}^{(n)}/\epsilon_{\mathrm{BB}}^{(n)}$ with respect to the ratio of the number of A loci to that of B loci, $N_{\mathrm{A}}^{(n)}/N_{\mathrm{B}}^{(n)}$. Note that $\epsilon_{\mathrm{AA}}^{(n)}/\epsilon_{\mathrm{BB}}^{(n)}$ is closely related to how A and B loci are distributed along a given chromosome rather than how many A and B loci exist.

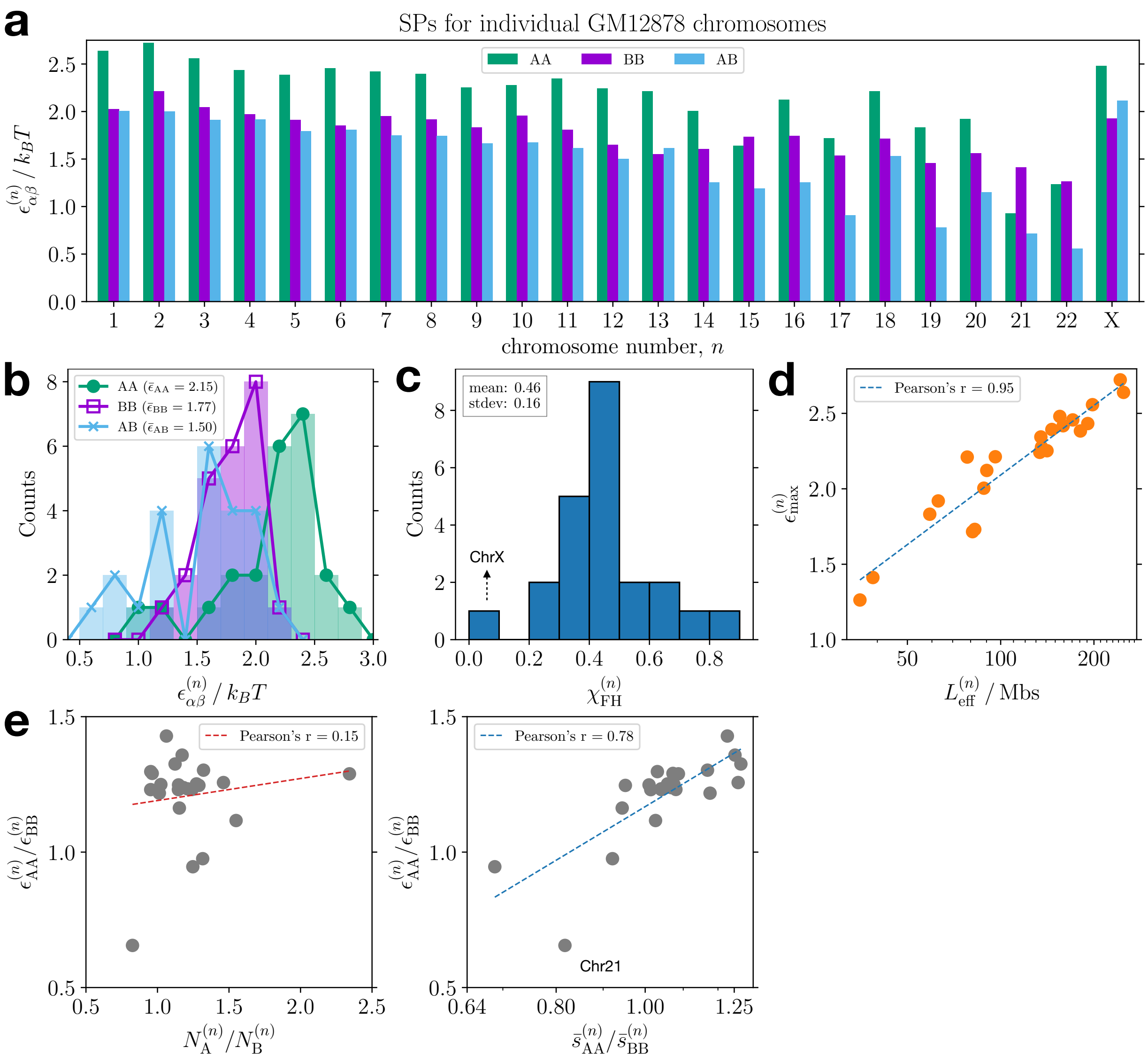


FIG. S7. SP values for GM12878 chromosomes show a similar trend as for IMR90. (a) A bar graph showing the interaction parameters based on SP at 50-kb resolution for individual GM12878 chromosomes. (b) Histograms of the effective interaction parameters shown in the bar graph of panel a, where the mean value for each pair type is given in units of $k_BT$ in the legend. (c) Distribution of the effective Flory-Huggins $\chi$ parameter computed from the extracted interaction energies, where the dotted arrow indicates the data point for ChrX. (d) Scatter plot showing the correlation between the maximum value of the interaction parameter for a given chromosome and its effective length along with its linear fit given by the dashed line, where the $x$-axis is shown in a log scale. (e) Scatter plots of the ratio between $\epsilon_{AA}$ and $\epsilon_{BB}$ for each chromosome with respect to the ratio of the number of A loci to that of B loci (left), and the ratio of average genomic distance for A-A pairs to that for B-B pairs (right). For the plot in the right, the $x$-axis is shown in the log scale.

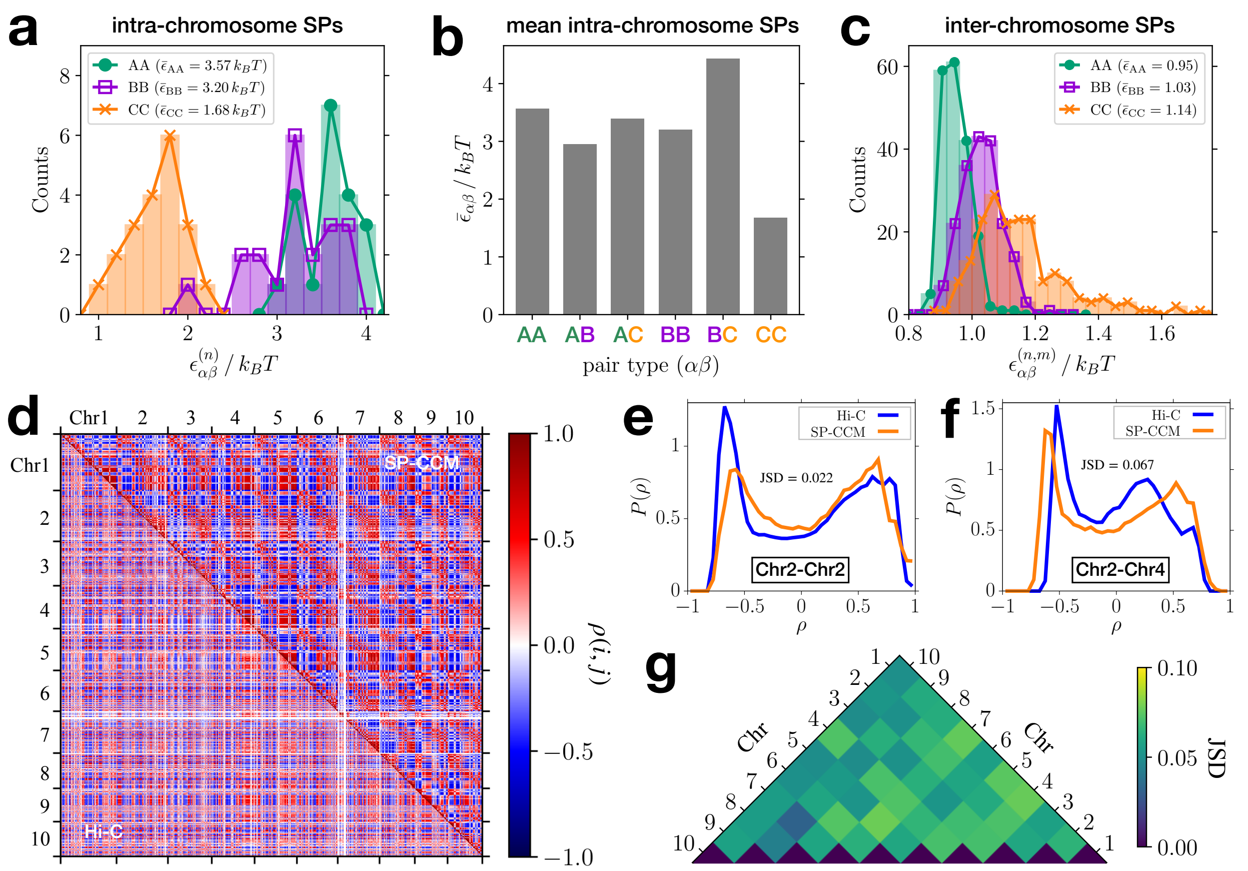


FIG. S8. SPs for inverted nuclei and SP-CCM simulation results. (a) Distributions of the SP-based energetic parameters for A-A, B-B, and C-C interactions for a given chromosome, extracted from the Hi-C CM for the inverted nuclei [28]. (b) Bar graph showing the average of the intra-chromosome values of the interaction energies for each pair type. (c) Histograms of the SP-based energetic parameters for A-A, B-B, and C-C interactions between different chromosomes, $\epsilon_{\alpha\beta}^{(n,m)}$ [Eqs. (17)-(19)]. The average for each pair type is given in the legend. (d) Pearson correlation matrices corresponding to the CMs for Chr1 to Chr10 shown in Fig. 4**e** in the main text. (e, f) Probability distributions of the Pearson correlation coefficients in Figs. 4**f** (left) and 4**g** (right), computed from the Hi-C (blue) and the SP-CCM simulations (orange). (g) Heat map showing the JSD value between the experimental and simulated distributions of the Pearson correlation coefficients for each chromosome pair. The overall mean JSD value is 0.061, which shows that the agreement between the distributions is excellent.

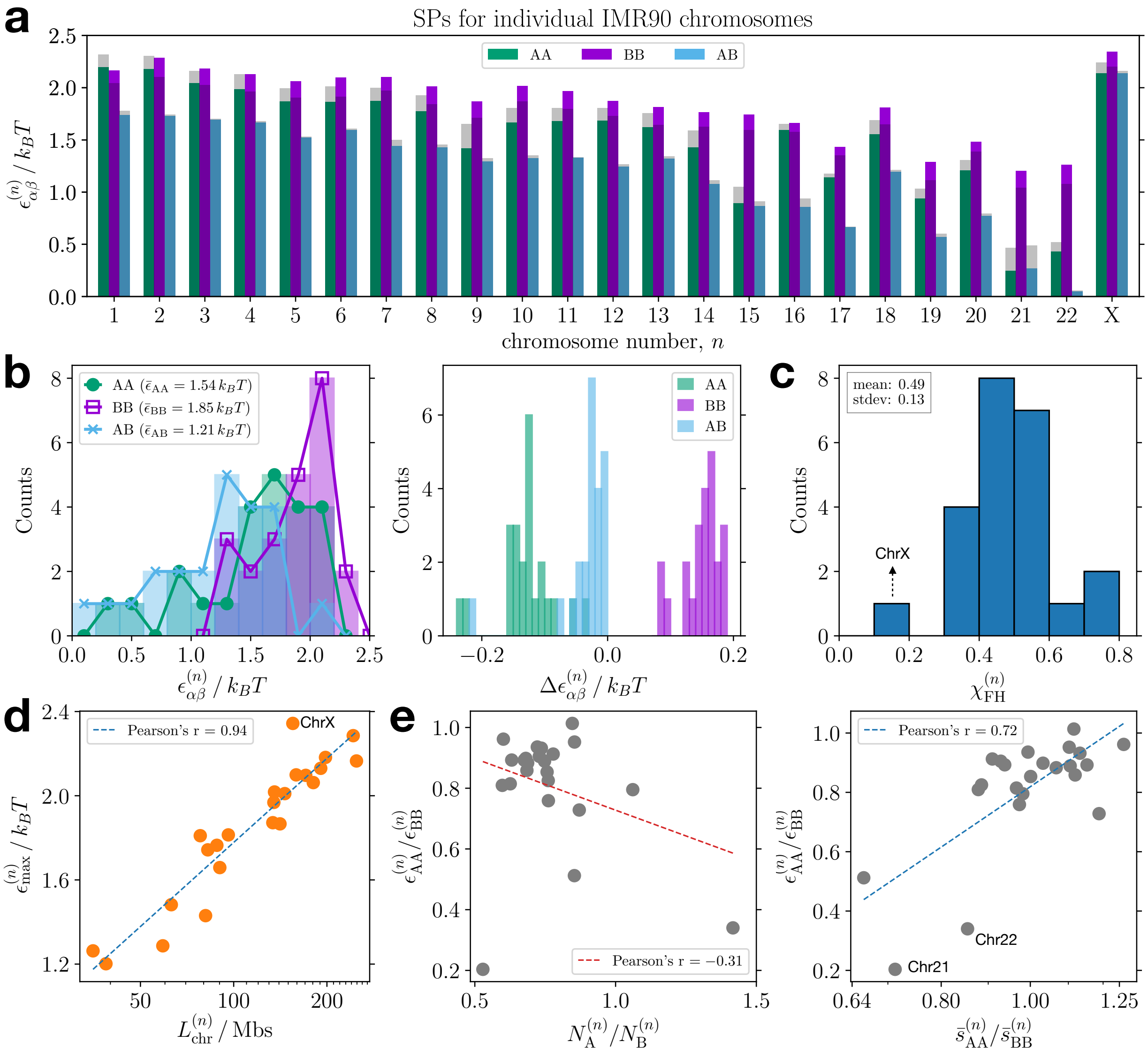


FIG. S9. Normalization of Hi-C contact maps with a matrix balancing method does not affect the trend in SPs significantly. (a) A bar graph showing the SP-based interaction parameters for individual IMR90 chromosomes, $\epsilon_{\alpha\beta}^{(n)}$, inferred from the normalized Hi-C contact map at 50-kb resolution [44]. The shaded bars indicate the values of $\epsilon_{\alpha\beta}^{(n)}$ inferred from the raw Hi-C data. (b) Histograms of $\epsilon_{\alpha\beta}^{(n)}$ obtained from the normalized Hi-C data, with the mean value for each pair type shown in the legend (left), and the difference between $\epsilon_{\alpha\beta}^{(n)}$ values from the normalized and raw Hi-C data (right). (c) Distribution of the effective Flory-Huggins $\chi$ parameter calculated from $\epsilon_{\alpha\beta}^{(n)}$ based on the normalized Hi-C data. (d) Scatter plot showing the correlation between the maximum value of $\epsilon_{\alpha\beta}^{(n)}$ for a given chromosome and the effective length along with the linear fit given by the dashed line, where the $x$-axis is shown in the log scale. (e) Scatter plots of the ratio between $\epsilon_{AA}$ and $\epsilon_{BB}$ for each chromosome with respect to the ratio of the number of A loci to that of B loci (left) and the ratio of average genomic distance for A-A pairs to that for B-B pairs (right). For the plot on the right, the $x$-axis is shown in a log scale.

| $n$ | $\epsilon^{(n)}_{\mathrm{AA}}/k_BT$ | $\epsilon^{(n)}_{\mathrm{BB}}/k_BT$ | $\epsilon^{(n)}_{\mathrm{AB}}/k_BT$ | $\chi^{(n)}_{\mathrm{FH}}$ |
|---|---|---|---|---|
| 1 | 2.31 | 2.04 | 1.78 | 0.40 |
| 2 | 2.30 | 2.10 | 1.74 | 0.46 |
| 3 | 2.16 | 2.02 | 1.70 | 0.39 |
| 4 | 2.13 | 1.96 | 1.68 | 0.37 |
| 5 | 2.00 | 1.91 | 1.53 | 0.42 |
| 6 | 2.01 | 1.91 | 1.61 | 0.36 |
| 7 | 2.00 | 1.97 | 1.50 | 0.48 |
| 8 | 1.93 | 1.84 | 1.45 | 0.43 |
| 9 | 1.65 | 1.71 | 1.32 | 0.36 |
| 10 | 1.81 | 1.87 | 1.35 | 0.48 |
| 11 | 1.80 | 1.79 | 1.33 | 0.47 |
| 12 | 1.81 | 1.73 | 1.26 | 0.50 |
| 13 | 1.75 | 1.64 | 1.34 | 0.36 |
| 14 | 1.59 | 1.63 | 1.11 | 0.49 |
| 15 | 1.05 | 1.59 | 0.91 | 0.41 |
| 16 | 1.65 | 1.57 | 0.94 | 0.68 |
| 17 | 1.18 | 1.35 | 0.67 | 0.60 |
| 18 | 1.69 | 1.65 | 1.21 | 0.46 |
| 19 | 1.03 | 1.11 | 0.60 | 0.47 |
| 20 | 1.31 | 1.39 | 0.79 | 0.55 |
| 21 | 0.46 | 1.04 | 0.49 | 0.26 |
| 22 | 0.52 | 1.07 | 0.06 | 0.74 |
| X | 2.24 | 2.20 | 2.16 | 0.06 |

TABLE S1. List of the mean SP values, $\epsilon^{(n)}_{\mathrm{AA}}$, $\epsilon^{(n)}_{\mathrm{BB}}$, and $\epsilon^{(n)}_{\mathrm{AB}}$, extracted from the Hi-C data for IMR90 chromosomes at 50-kb resolution and the corresponding Flory-Huggins parameter, $\chi^{(n)}_{\mathrm{FH}}$.
